# Disparate regulation of *imd* drives sex differences in infection pathology in *Drosophila melanogaster*

**DOI:** 10.1101/2020.08.12.247965

**Authors:** Crystal M. Vincent, Marc S. Dionne

## Abstract

Male and female animals exhibit differences in infection outcomes. One possible source of sexually dimorphic immunity is sex-specific costs of immune activity or pathology, but little is known about the independent effects of immune-induced versus microbe-induced pathology, and whether these may differ for the sexes. Here, through measuring metabolic and physiological outputs in wild-type and immune-compromised *Drosophila melanogaster*, we test whether the sexes are differentially impacted by these various sources of pathology and identify a critical regulator of this difference. We find that the sexes exhibit differential immune activity but similar bacteria-derived metabolic pathology. We show that female-specific immune-inducible expression of *PGRP-LB*, a negative regulator of the Imd pathway, enables females to reduce immune activity in response to reductions in bacterial numbers. In the absence of *PGRP-LB*, females are more resistant of infection, confirming the functional importance of this regulation and suggesting that female-biased immune restriction comes at a cost.

## Introduction

Biological sex can influence an animal’s response to infection, with females often mounting stronger innate and adaptive immune responses compared to males. Across multiple taxa, the sexes exhibit differing incidences of infection, pathogen loads, pathogen-derived virulence, and immune efficacy [1–8]. In humans, the greater responsiveness of the female immune response can confer rapid pathogen clearance, reduced mortality rates and greater efficacy of vaccines; however, it also thought to be responsible for the increased incidence of inflammatory and autoimmune disease in women [3,9,10]. Thus, females appear to trade-off the rapid and efficient clearance of foreign bodies, with the risk of doing self-harm, either due to autoimmunity or immunopathology. Consequently, sex-specific infection outcomes could be driven by differences between the sexes in the risks of autoimmunity, immunopathology, virulence (pathogen-induced harm), or trade-offs between immunity and other important traits.

The origins of infection-induced pathology, and the mechanisms employed by the host to limit pathology, are key issues in understanding this difference between the sexes. Infection pathology can result from direct interactions between host and pathogen, or can be driven indirectly. Direct pathology is caused by the pathogen itself and its products. It can be produced by many effects; pathogen- or pathogen effector-driven damage to host tissue [11,12] is the most obvious of these, but other direct pathological processes include competition with the host for access to resources [13–15]. Indirect pathology in contrast, is caused not by the pathogen itself, but by some aspect of the host response to pathogen. This is most often conceived as pathology caused by immune effectors; other indirect pathologies come in the form of immune trade-offs, where immune activation leads to the reallocation of host resources from other processes, such as longevity, reproduction, competitive ability and development [5,16–23].

Differences in infection outcomes between hosts can result from differences in the ability of the host to clear the pathogen (“resistance” mechanisms) or from differences in sensitivity to direct or indirect pathology (“tolerance” mechanisms). In any given infection, the survival and continued health of the host will be the product of a complex interaction of host and pathogen genotype as well as other circumstances. It is unclear whether the well-documented effects of host sex on infection outcome in general primarily originate in changes in resistance to the infectious agent or in tolerance of direct or indirect pathology.

To distinguish these effects, we used the fruit fly, *Drosophila melanogaster* and consider the response of wild-type and immunocompromised flies to infection with the bacterium *Escherichia coli.* Unlike mammals, *D. melanogaster* lacks an adaptive immune response, instead, flies have a well-developed innate immune response consisting of both cellular and humoral components. The humoral response of *D. melanogaster* involves the inducible production of circulating factors – primarily antimicrobial peptides (AMPs) – that are directly microbicidal. Though infection with *E. coli* is non-lethal and efficiently controlled by the immune response of wild-type flies, *E. coli* infection cannot be controlled in immunocompromised flies [24]. Therefore, using this system, we sought to distinguish between pathology resulting from the immune response and pathology resulting from the microbe. We test whether the sexes are differentially impacted by these two sources of pathology using multiple metabolic and physiological measures as readouts. We show that females reduce the costs of immune activity via strict regulation of the Imd pathway and that this comes at the cost of bacterial clearance.

## Results

### Male and female flies exhibit differences in IMD pathway function after infection

To determine whether male and female flies exhibited any significant difference in their ability to defend against non-pathogenic Gram-negative bacterial infection, we first measured survival and bacterial numbers after infection with *E coli* of wild-type flies. Prior work in our lab as well as others has found that *D. melanogaster* infected with *E. coli* either eliminate the bacteria or maintain them at low levels at no obvious cost to the host [25,26]. As expected, we did not find a strong effect of infection with live or dead (heat-killed) *E. coli* on the lifespan of wild-type flies (Fig. 1A; S1 Fig.; S1 Table). However, when we compared bacterial numbers between infected males and females, we found a clear trend toward larger numbers of surviving bacteria in females; this trend became significantly different 6 hours after infection (Fig. 1B).

**Figure 1.**
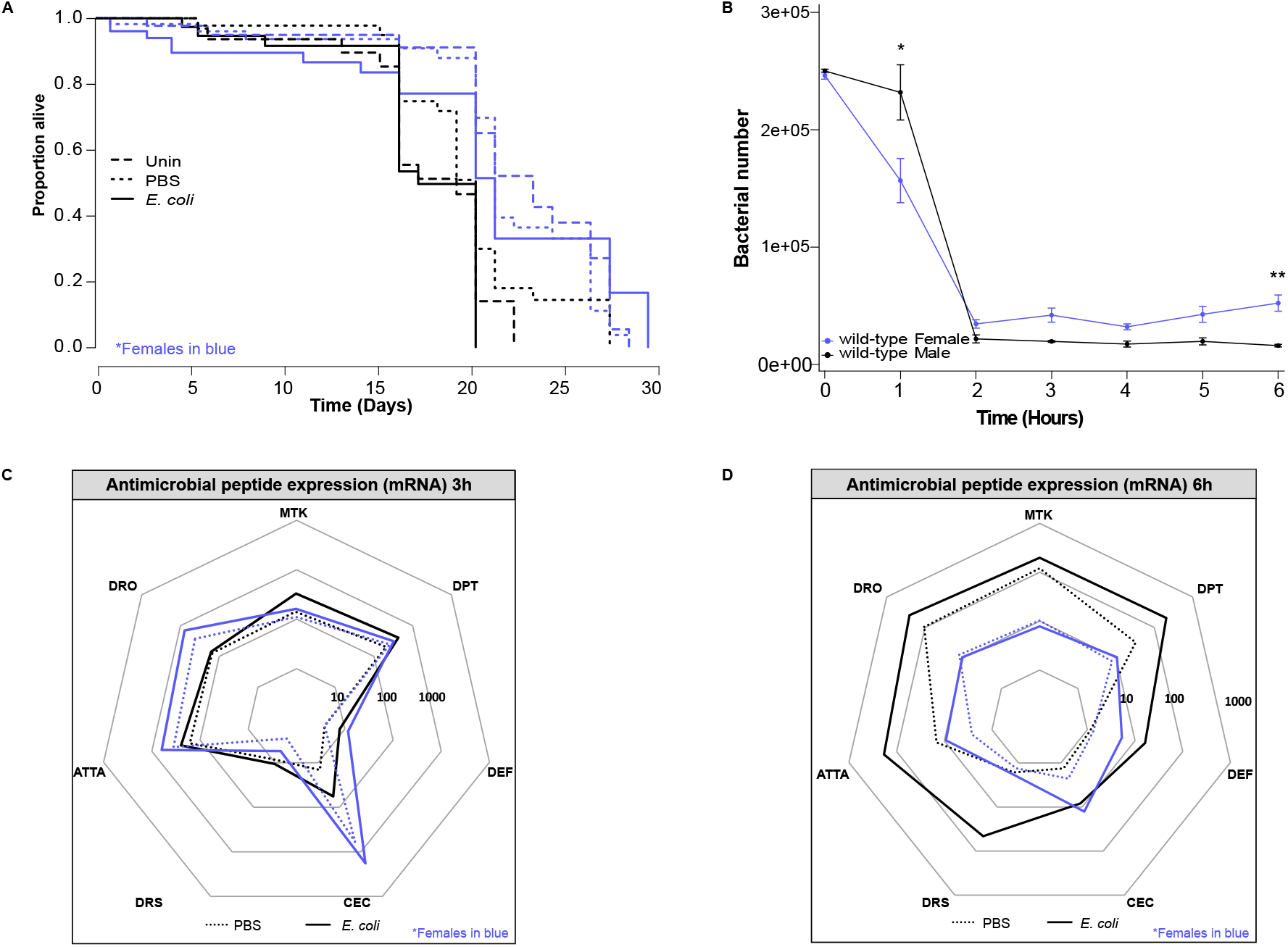
Sex-specific outcomes of *E. coli* infection in wild-type flies. Representation in all plots: males - black; females - blue. **(A)** Survival of *E. coli* infected flies. *Escherichia coli* infected flies are indicated by solid lines. Uninfected and PBS controls are indicated by long and short dashed lines, respectively. Flies had an average median survival across all treatments of 21.5d and 18.5d for females and males, respectively (Coxph: df = 7, n = 396, Wald test = 43.75, p = 2e-07). There was no effect of treatment on survival in either sex. Survivals were performed at least twice, each repeat included 20-40 flies/treatment. **(B)** Bacterial quantification in wild-type flies. Wild-type females had fewer bacteria than males 1h after injection (t-test = −2.495, p = 2.03e-02, n = 26) but more bacteria at 6h (t-test = 5.397, p = 1.3e-03, n = 25). No significant difference in bacterial load in wild-type flies was observed at any other time. Markers indicate means and bars represent SE. Statistical significance: * p<0.05; ** p<0.01. Quantifications were performed twice, each repeat included 6-8 biological replicates consisting of 1 fly each. **(C and D)** Antimicrobial peptide transcript levels 3h (C) and 6h (D) post infection in wild-type flies. Expression is shown relative to uninfected flies of the same genotype/sex. On average, infected males had AMP transcript levels 16x greater than females (*Mtk-*25x; *DptA*-19x; *Def*-3x; *CecA1*-0.65x; *Drs*-25x; *AttaA*-19x; *Dro*-23x). Solid lines represent infection with *E. coli* whilst dotted are PBS injected. The area contained within the innermost heptagon represents induction levels falling between one and ten times that of the uninfected controls. The middle and outer heptagons represent 100 and 1000-fold induction, respectively. These data are also shown, represented differently, in S1 Fig. AMP assays were performed 2 - 4 times, each repeat included 3 or 4 biological replicates/treatment consisting of 3 flies each.

**Table 1.**
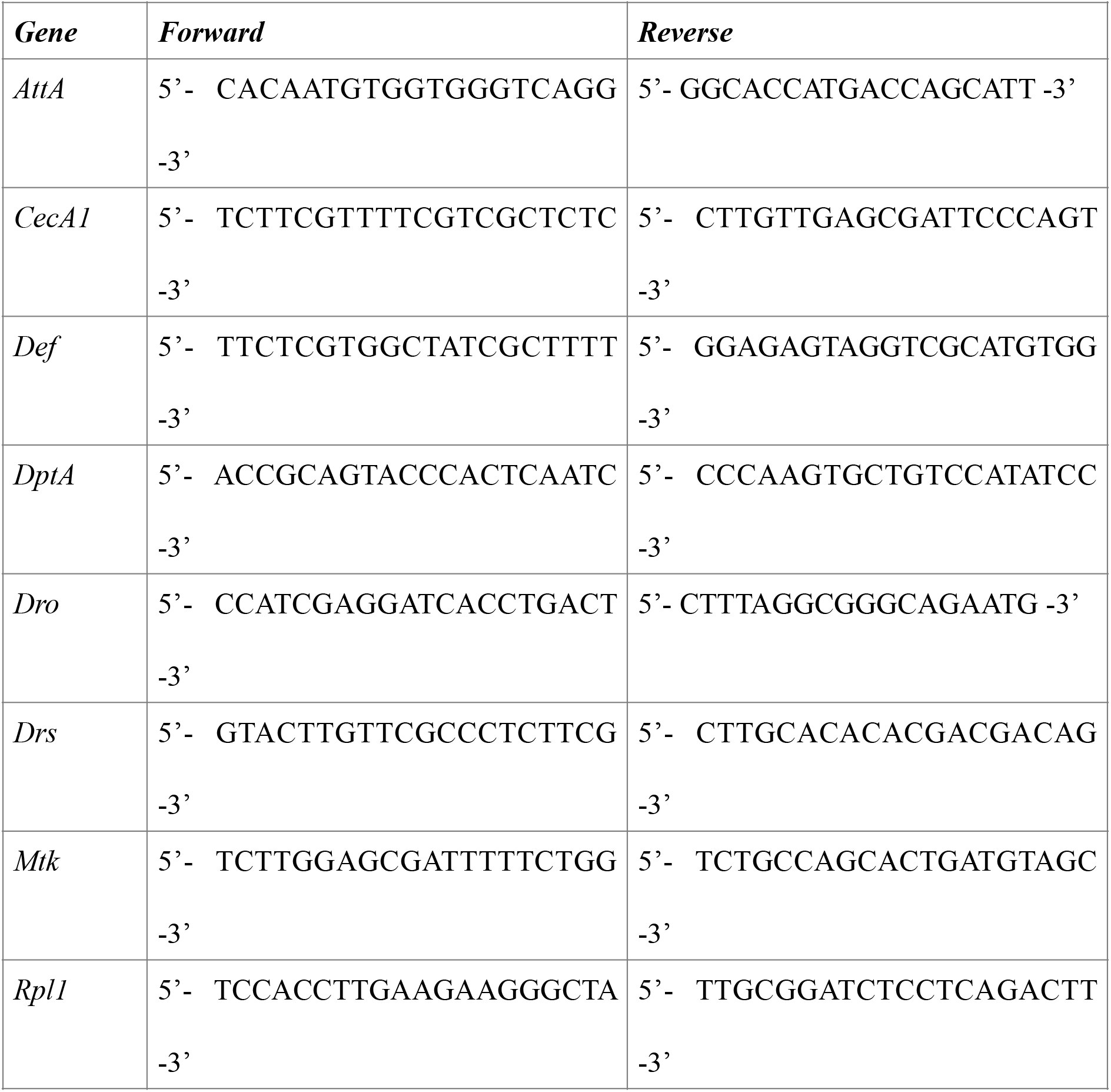
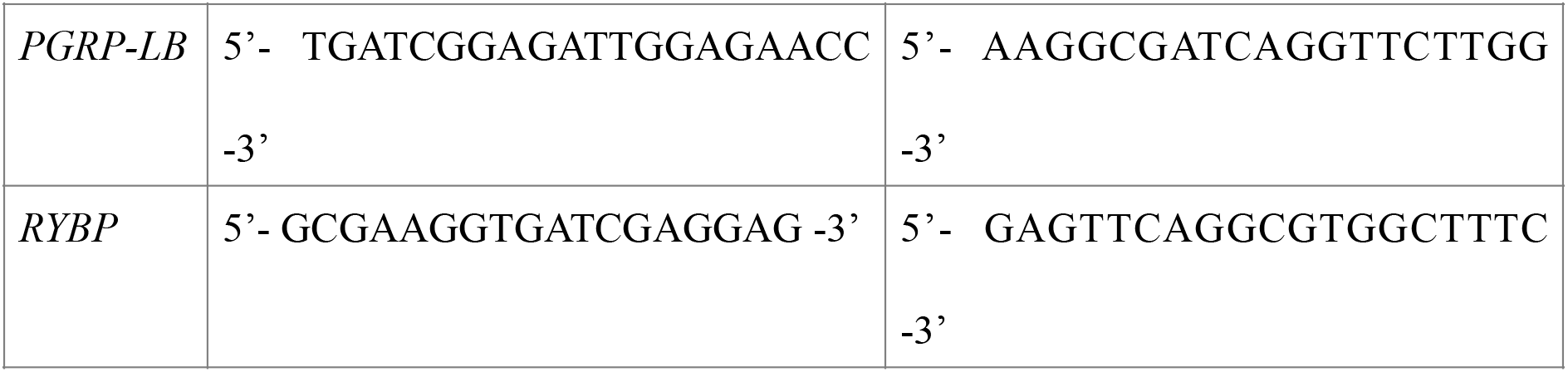
Primer sequences used for qRT-PCR.

Defence against *E coli* infection is expected to depend primarily on the activity of the IMD signalling pathway and its antimicrobial peptide target genes [27,28]. The fact that males and females exhibited differences in bacterial numbers led us to examine antimicrobial peptide mRNA expression 3 and 6 hours after infection; these times were chosen because 3 hours was not long after the bulk of bacterial killing had been achieved, while 6 hours is the reported peak of *Diptericin* induction – a canonical read-out of *imd* activity – in wild-type animals [29]. At 3 hours after infection, male and female flies exhibited broadly similar levels of antimicrobial peptide transcripts (Fig. 1C; S1 Fig.). However, by 6 hours after infection, antimicrobial peptide expression was significantly reduced in female flies relative to males, despite females having higher bacterial numbers (Fig. 1D; S1 Fig.). Importantly, AMP levels were notably greater in infected females 3h following injection than they were at 6h, whilst male levels were unchanged, suggesting that females were more responsive to bacterial load as a cue to shut down immune activity.

### Loss of *imd* reveals a sex-specific tolerance to *E. coli* infection

The fact that we found the regulation of IMD signalling was different between the sexes, led us to look more closely at the sex-specific consequences of the loss of *imd* function during *E coli* infection. We infected *imd* mutants with *E. coli*; we also injected a subset of these flies with latex beads to inhibit their ability to phagocytose bacteria [30], resulting in flies with both phagocytosis and antimicrobial peptide activity inhibited. As expected, *imd* mutants of both sexes had significantly reduced survival when infected with *E. coli* compared to their PBS and uninfected controls (Fig. 2A). However, this effect was particularly strong in males: male *imd* mutants lived roughly 60% as long as females. Inhibiting the phagocytic response with latex beads did not affect survival in either sex (S2 Fig.), further supporting the idea that AMP activity plays the primary role in this infection. When we examined bacterial loads in male and female *imd* mutants, we found that both sexes carried identical numbers of bacteria, indicating that the difference in survival between male and female animals reflected different levels of infection tolerance (Fig. 2B). The fact that this differential tolerance effect was revealed only in *imd* mutants implied that it was a consequence of different degrees of non-IMD pathway immune activation—that the secondary immune response pathways revealed by *imd* mutation were more damaging to males, possibly because of quantitative differences in their activation between the sexes.

**Figure 2.**
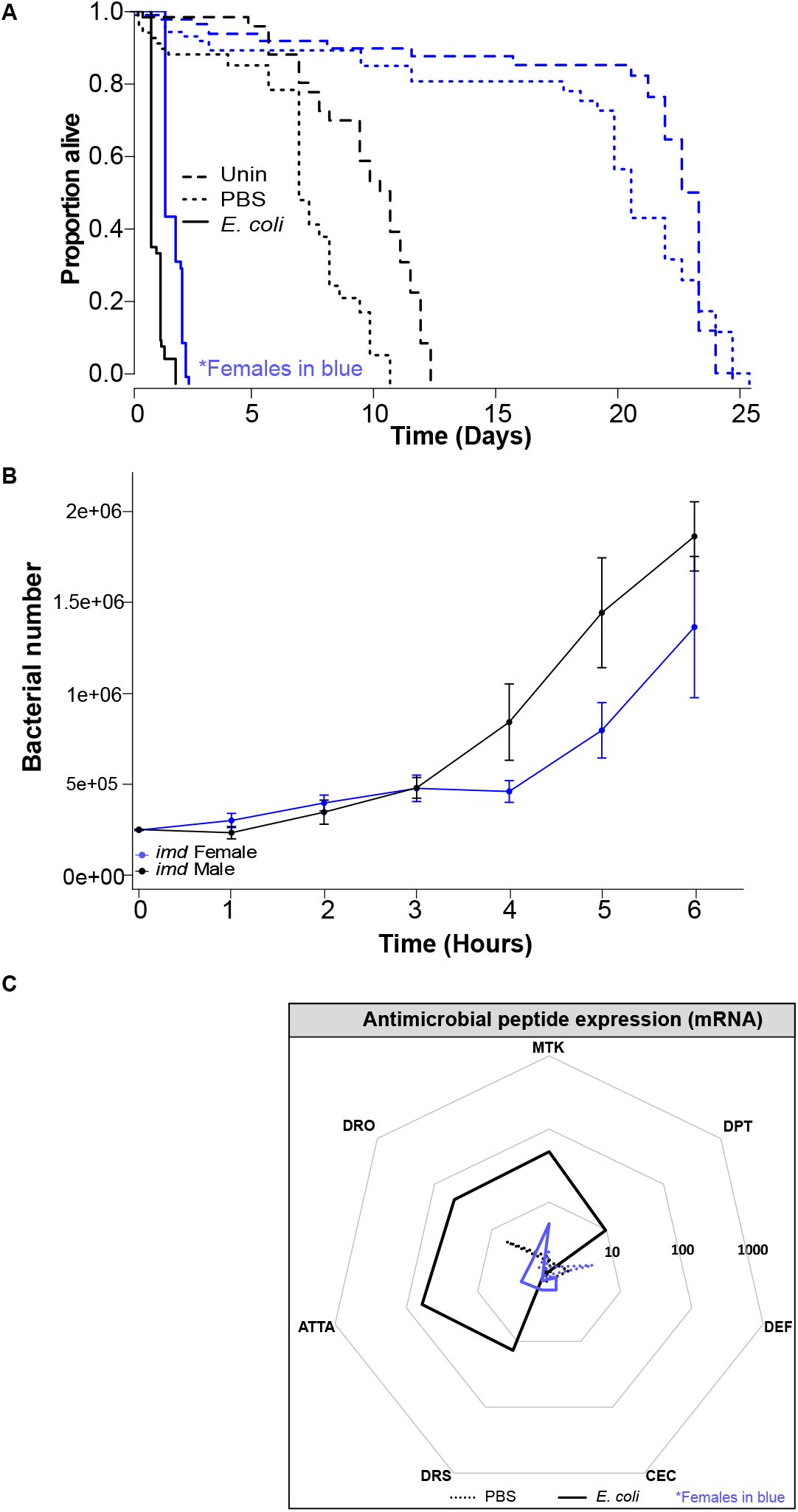
Sex-specific outcomes of *E. coli* infection in *imd* flies. Representation in all plots: males - black; females - blue. **(A)** Survival of *E. coli* infected flies. *Escherichia coli* infected flies are indicated by solid lines. Uninfected and PBS controls are indicated by long and short dashed lines, respectively. Median survival of *E. coli* infected *imd* flies was 41h and 27h for females and males, respectively (Coxph: df = 9, n = 255, Wald test = 126.2, p = <2e-16). Both PBS and infection reduced male survival whilst female survival was only affected by infection. Survivals were performed at least twice, each repeat included 20-40 flies/treatment. **(B)** Bacterial quantification in *imd* mutant flies. Flies exhibited no significant difference in bacterial number at any time. Markers indicate means and bars represent SE. Quantifications were performed twice, each repeat included 6-8 biological replicates consisting of 1 fly each. **(C)** Antimicrobial peptide transcript levels 6h post infection in *imd* mutant flies. Expression is shown relative to uninfected flies of the same genotype/sex. Solid lines represent infection with *E. coli* whilst dotted are PBS injected. The area contained within the innermost heptagon represents induction levels falling between one and ten times that of the uninfected controls. The middle and outer heptagons represent 100 and 1000-fold induction, respectively. These data are also shown, represented differently, in S2 Fig. AMP assays were performed 2 - 4 times, each repeat included 3 or 4 biological replicates/treatment consisting of 3 flies each.

We tested this possibility by assaying antimicrobial peptide induction in *imd* mutants infected with *E coli*. Females exhibited no response at all, while males exhibited a residual 10-100 fold induction of most antimicrobial peptides (Fig. 2C; S2 Fig.). This level of expression was clearly insufficient for antimicrobial activity, as the sexes exhibited identical bacterial numbers, but was a potential cause of pathology in *imd* mutant males.

### Infection with *E. coli* leads to depletion of triglycerides

Because resources are finite, individuals must manage investments in multiple biological processes. The ability to draw on metabolic reserves of triglyceride or glycogen allows animals to run temporary metabolic deficits in response to unexpected costs (e.g. immunity). We hypothesised that the sex differences we observed in immune activity and tolerance of infection in wild-type and *imd* mutant flies might also be reflected in differences in the metabolic cost of infection. To test this, we assayed levels of free sugar (glucose and trehalose), stored carbohydrate (glycogen), stored triglyceride, and respiration in wild-type and *imd* flies. Previous studies in *D. melanogaster* found that lethal bacterial infections can lead to hyperglycaemia, as well as a reduction in triglyceride and glycogen stores, but these metabolites had not been examined during acute infection with nonpathogens [31–33].

We found that 6h post infection with *E. coli*, wild-type flies had significantly less stored triglyceride than their PBS controls; this effect was independent of both genotype and sex (Fig. 3A; S2 Table). Importantly, infection with heat-killed *E. coli* did not deplete triglyceride, indicating that this effect is pathogen-driven. Wild-type males had significantly less circulating sugars but more glycogen stores than females, but neither of these was changed by infection. Respiration was unaffected by infection status in wild-type flies (S3 Fig.). *imd* mutants exhibited a somewhat different pattern to wild-type. While there was no effect of infection on glycogen, free sugar levels appeared to be elevated following wounding and infection in males, this increase was not significant, possibly due to its extreme variability; this effect on free sugar was absent in females (Fig. 3B). As in wild-type flies, both male and female *imd* mutants exhibited significant reduction in triglyceride resulting from infection and this effect was notably stronger in males (Fig. 3B; S2 Table). This fit with our observation that *imd* mutant males exhibited a stronger (though clearly ineffective) immune response to *E coli* infection than *imd* mutant females, as a possible cause for greater triglyceride depletion in males could be increased demands resulting from immune activity.

**Figure 3.**
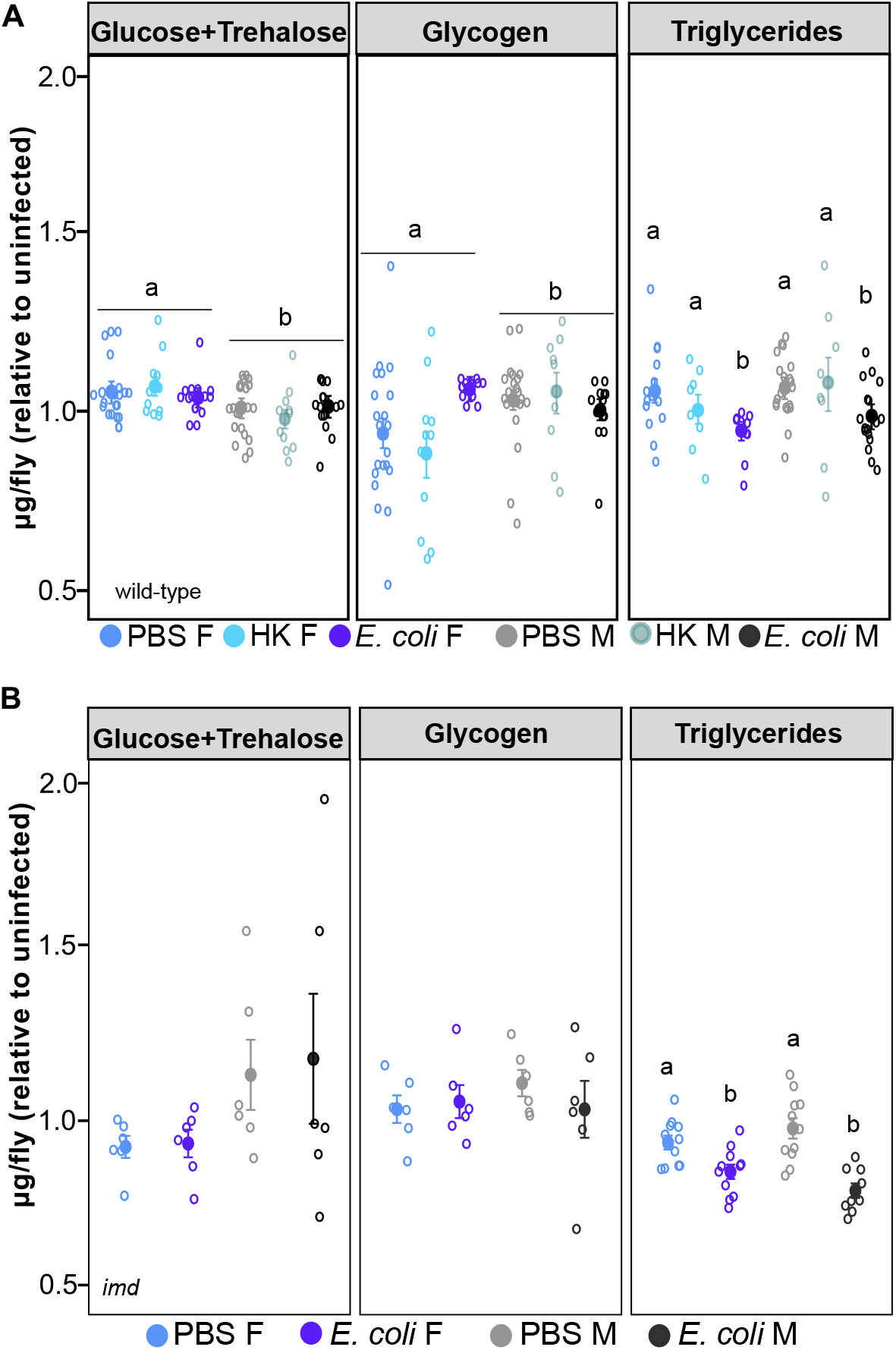
No sex difference in metabolic pathology of *E. coli* infection in wild-type and *imd* flies. Triglyceride and carbohydrate levels in live and heat-killed *E. coli-*infected flies. **(A)** In wild-type there was an effect of sex on circulating sugar (AOV: df = 1, n = 67, F =14.7, p = 2.7e-04) and glycogen levels (AOV: df = 1, n = 59, F = 6.15, p = 0.016) with males having less circulating sugar but more glycogen stores than females. There was also an effect of infection status on triglyceride levels, such that *E. coli* infection led to triglyceride loss (AOV: df = 2, n = 74, F = 5.73, p = 4.8e-02). There was no interaction between sex and infection on triglyceride loss. **(B)** *imd* flies showed no effect of sex nor infection status on circulating sugar and glycogen levels. There was no effect of sex on triglyceride levels, but there was an overall effect of both treatment (AOV: df = 1, n = 49, F = 44.971, p = 2.8e-08) and the interaction between sex and treatment on triglycerides; *E. coli* infection led to triglyceride depletion in both sexes, relative to their PBS controls (AOV: df = 1, n = 49, F = 7.417, p = 9.2e-03; M PBS-M *E. coli*, p-adjusted = 4.0e-07; F PBS-F *E. coli* p-adjusted = 1.01e-02). Bars indicate SE. Letters indicate statistical groupings. Full statistics including non-significant results can be found in S2 Table. All assays were performed 2 or 3 times, each repeat included 4 biological replicates/treatment consisting of 3 (carbohydrates) or 8 (triglycerides) flies each.

Because animals spend significant energy on reproduction, and reproductive effort is likely to restrict or trade-off with immunity [34], we assayed reproductive output during infection. We placed infected flies in tubes with flies of the opposite sex and ‘competitors’ of the same sex, but of a different genotype (*Dh44*[3xP3-DsRed]). We allowed flies to mate for 12h and then discarded adults. Offspring resulting from matings with competitors were easily identifiable by their red-fluorescent eyes. Males were less likely to have a successful mating interaction than females, but neither sex showed an effect of infection on mating success or the number of offspring produced (S4 Fig.). These findings demonstrate that despite observing metabolic shifts and sex-specific AMP induction and pathology (bacterial load), reproductive output is unaffected in the short term by *E. coli* infection.

### Sex-specific expression of IMD pathway regulators

We have shown that male and female flies exhibit clear differences in the dynamics of the transcriptional response to *E coli* infection, presumably due to distinct mechanisms of immune regulation, and that in flies lacking the IMD pathway male animals exhibit distinctly greater responses to infection in terms of gene expression and triglyceride depletion and die more rapidly than females. We wished to gain some mechanistic insight into these differences between the sexes, so we analysed the expression of known negative regulators of IMD signalling in male and female flies. We expected that negative regulators responsible for the effects we observed on antimicrobial peptide expression should be more inducible in females.

Several negative regulators of Imd pathway activity have been described [35–38]. We assayed several of these regulators for increased infection inducibility in female flies relative to males (S5 Fig.). Two negative regulators – *PGRP-LB* and *RYBP* – were expressed at higher levels specifically in *E. coli* infected females 3 hours post-infection (Fig. 4A, B). A more detailed analysis of the timecourse of expression of *PGRP-LB* and *RYBP* revealed that both were upregulated as early as one hour after infection in females, and both showed continuing strong expression 3 hours after infection, especially in females (Fig. 4A, B; S5 Fig.). However, by 6 hours after infection, PGRP-LB expression had returned to near-normal in both males and females, while RYBP expression was now induced in males to the same high level seen from 1 hour in females. This difference in the regulatory timing of the Imd pathway can be seen when we compare AMP expression at 3 and 6h in each sex (Fig. 4C, D).

**Figure 4.**
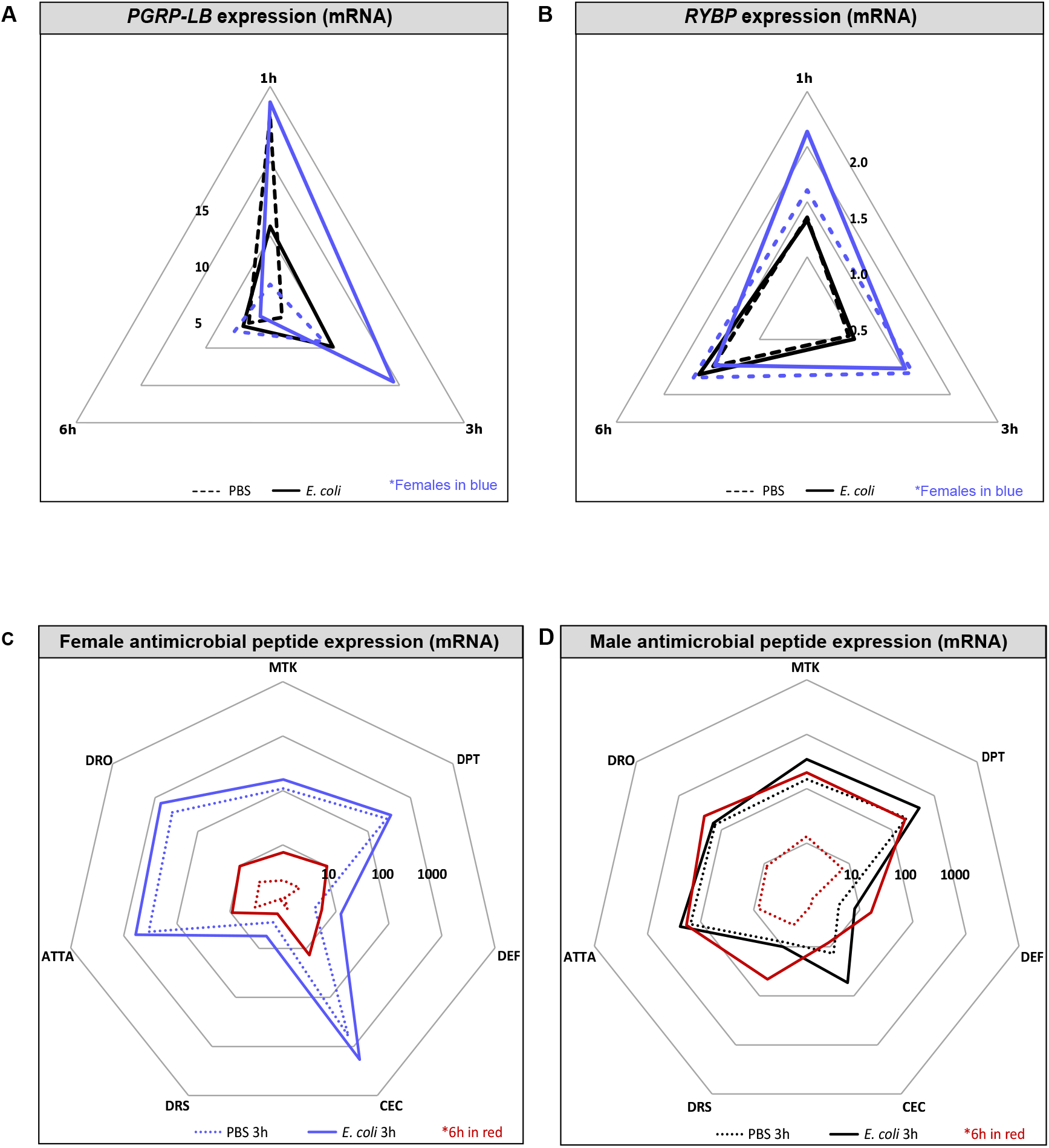
Sex-specific temporal regulation of *imd* during *E. coli* infection wild-type flies. Representation in all plots: males - black; females - blue. **(A and B)** Expression of *PGRP-LB* (A) and *RYBP* (B) 1, 3, and 6 hours after infection in male and female flies. Plotted values are relative to the uninfected controls. Solid lines represent infection with *E. coli* whilst dotted lines represent PBS injection. These data are also shown, represented differently, in S5 Fig. **(C and D)** Antimicrobial peptide transcript levels in females (C) and males (D) 3h and 6h after infection. Expression is shown relative to uninfected flies of the same genotype/sex. Solid lines represent infection with *E. coli* whilst dotted are PBS injected. Data collected at 6h are indicated in red. The area contained within the innermost heptagon represents induction levels falling between one and ten times that of the uninfected controls (downregulation was not observed in any of the tested genes). The middle and outer heptagons represent 100 and 1000-fold induction, respectively. AMP assays were performed 2 - 4 times, each repeat included 3 or 4 biological replicates/treatment consisting of 3 flies each.

*PGRP-LB* is an amidase that degrades the DAP-type peptidoglycan of Gram-negative bacteria, dampening activation of the Imd pathway by degrading the activating ligand [38]. In contrast, *RYBP* inhibits Imd pathway activity by promoting proteasomal degradation of the pathway’s NF-κB transcription factor, *Relish* [36]. *PGRP-LB* was of particular interest because it reduces pathway activity by degrading free peptidoglycan – that is, it will reduce pathway activity when the immune response has been effective; it was thus particularly interesting as a causal factor because rendering PGRP-LB infection-inducible would render the pathway responsive to its own success. A peptidoglycan-degrading activity also could regulate IMD-independent immune activities, which could explain the sex differences we observed in immune activity, metabolic impact, and infection pathology in *imd* mutants. We thus decided to analyse immune function in male and female *PGRP-LB* mutants.

### *PGRP-LB* mutants exhibit reversed sex bias in immunity, improved immune function and altered metabolic response to infection

To test whether *PGRP-LB* activity was responsible for the sex difference in immune function, we infected male and female *PGRP-LB* null mutants with *E. coli* and measured antimicrobial peptide expression, bacterial numbers, and survival of the host. In the absence of *PGRP-LB*, the male-biased AMP expression observed six hours following infection with *E. coli* was abolished (Fig. 5A). Moreover, these mutants had a strong and sex-specific effect on bacterial load. As in wild-type flies, *PGRP-LB^Δ^* mutants drastically reduced bacterial load within the first 2 hours post-infection, at which time bacterial numbers effectively plateaued. However, in contrast to what we saw in wild-type flies, *PGRP-LB^Δ^* males had a tendency to have more bacteria than females throughout the 6h period assayed (Fig. 5B), confirming our supposition that wild-type females downregulate AMP activity at a cost of resistance, and indicating that sex-specific *PGRP-LB* induction has important functional consequences for the realised immune response. The effect on overall lifespan was more complex: similar to what we observed in wild-type flies, uninfected *PGRP-LB^Δ^* females lived longer than males (Fig. 5C). Wounding had a significant impact on survival in both sexes, with both PBS and *E. coli* injected animals having reduced survival (though the two treatments did not differ from each other)., Because PGRP-LB should have little effect in the absence of peptidoglycan, the effect of sterile wounding was somewhat confusing; one possibility is that the previously-documented effect of PGRP-LB on interaction with microbiota-derived peptidoglycan may have specific importance in the regulation of immune responses following sterile injury [39].

**Figure 5.**
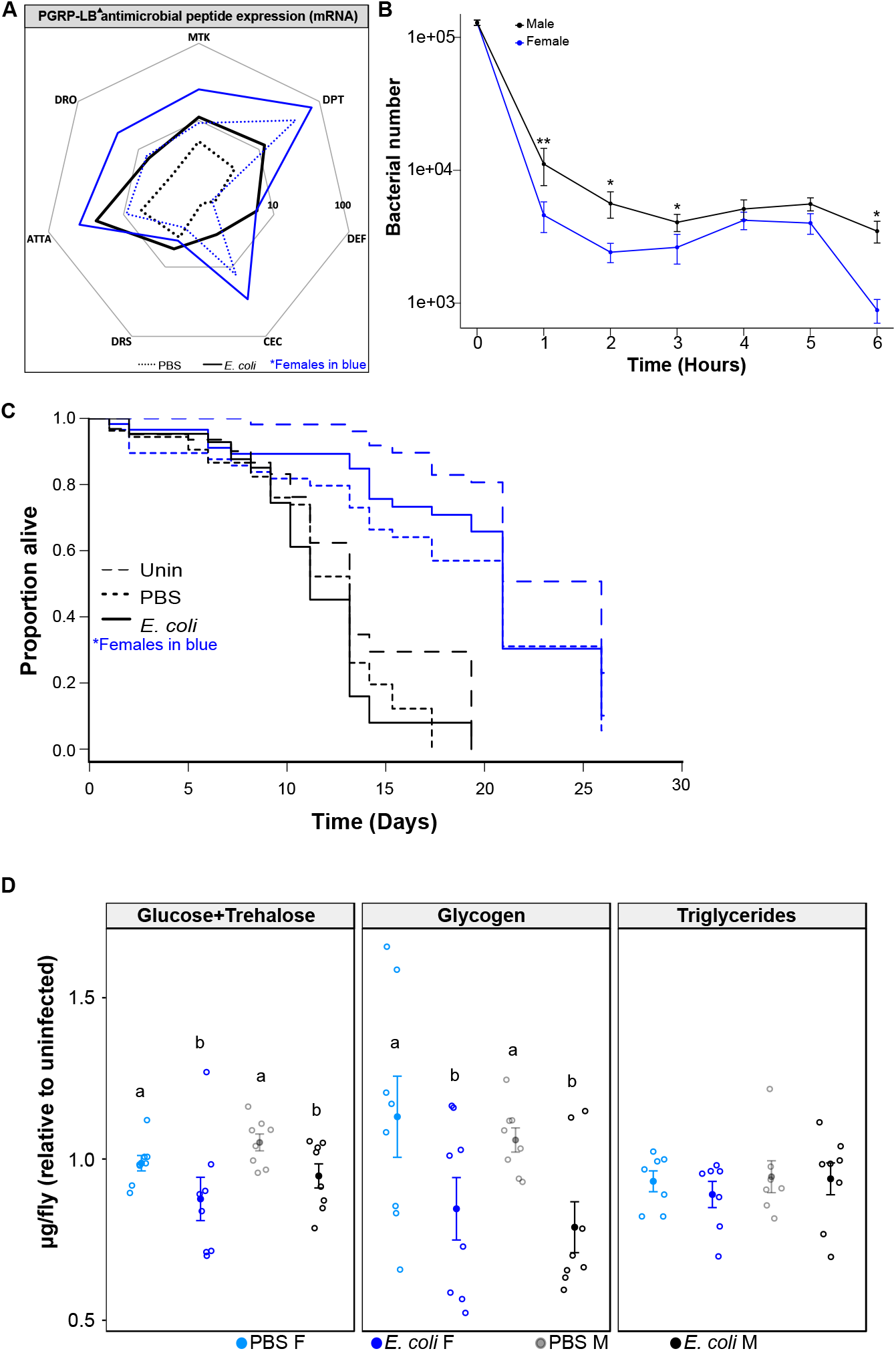
*PGRP-LB^Δ^* males and females exhibit parallel metabolic shifts during infection. Representation in all plots: males - black; females - blue. **(A)** AMP expression is shown relative to uninfected flies of the same genotype/sex. Solid lines represent infection with *E. coli* whilst dotted are PBS injected. The area contained within the innermost heptagon represents induction levels falling between one and ten times that of the uninfected controls. The outer heptagon represents 100-fold induction. Assays were performed twice, each repeat included 3-4 biological replicates/treatment consisting of 3 flies each. These data are also shown, represented differently, in S6 Fig. **(B)** Bacterial load observed over the first 6h of infection. Males had significantly higher bacterial loads throughout most of the observed period (1h: Mann-U = 21, p = 6.1e-03, n = 23; 2h Mann-U = 48, p = 0.039, n = 27; 3h: Mann-U = 52, p = 0.022, n = 29; 4h Mann-U = 75, p = 0.45, n = 27; 5h: t-test = −1.49, p = 0.15, n = 24; 6h Mann-U = 1.5, p = 0.012, n =12). Markers indicate means and bars represent SE. Statistical significance: * p<0.05; ** p<0.01. Quantifications were performed twice, each repeat included 8 biological replicates consisting of 1 fly each. **(C)** Survival of flies infected with *E. coli* indicated by solid lines. Uninfected and PBS controls are indicated by long and short dashed lines, respectively. *E. coli*-infected females had a median survival 86% greater than that of males (Female = 20.9d, Male = 11.2d; Coxph: df = 5, n = 344, Wald test = 125.5, p = 1.0e-16). Survivals were repeated twice, each repeat included 2 biological replicates/treatment consisting of 20-40 flies each. * Due to Covid-19, the second repeat was terminated early; unfortunately, this was done in a manner that precluded the assignment of treatment identity to surviving flies, as a result, these flies were removed from the experiment rather than censored. Thus, data after day 21 represent one experiment. * **(D)** Infection had a significant effect on circulating sugar such that the amount of circulating sugar in *E. coli*-infected animals was lower than in PBS controls (AOV: df = 1, n = 32, F = 6.44, p = 1.7e-02); whereas sex had no effect on circulating sugars, nor was there a significant interaction between the two. Similarly, *E. coli*-infection led to marked reduction in stored glycogen (AOV: df = 1, n = 32, F = 9.41, p = 4.8e-03), with no effect of sex, nor a significant interaction between sex and treatment. Neither infection status nor sex effected triglyceride levels. Large filled markers indicate means while smaller circles represent individual data points. Letters indicate statistical groupings. Bars indicate SE. All assays were performed twice, each repeat included 4 biological replicates/treatment consisting of 3 (carbohydrates) or 8 (triglycerides) flies each. Full statistics including non-significant results can be found in S3 Table.

We next aimed to identify the effects of *PGRP-LB* on the physiological consequences of immune activation—in particular, to explore the extent to which the metabolic consequences of acute infection are driven by host or pathogen-derived activities. We predicted that if triglyceride loss during *E. coli* infection in wild-type flies is driven entirely by pathogen-derived costs, that the nearly 10-fold reduction of bacterial load observed in infected *PGRP-LB^Δ^* flies might be sufficient to abrogate triglyceride loss; conversely, if triglyceride loss were driven by Imd pathway activity, the prolonged Imd pathway activation observed in *PGRP-LB* mutants should result in greater loss of triglyceride than in wild-type animals. We found that in *PGRP-LB^Δ^* flies, triglyceride levels were unaffected by *E. coli* infection, confirming that Imd pathway activity was not the cause of triglyceride depletion in this infection. Infected flies also had lower levels of circulating sugars and glycogen, independent of sex (Fig. 5D, S3 Table). This effect of infection on circulating and mobile energy observed in *PGRP-LB^Δ^* flies may be indicative of the energy requirement of a hyperactive immune response.

## Discussion

Differences between males and females in immune activity and infection outcomes are pervasive throughout the animal kingdom. Here, we have explored the differences between male and female *Drosophila* in their response to a non-pathogenic Gram-negative bacterial infection. Though both males and females could control this infection at the cost only of transient metabolic depletion, our analysis revealed that females maintained much stricter control of their own immune response; this was achieved by female-specific transcriptional induction of a peptidoglycan amidase that degrades peptidoglycan fragments liberated from bacteria after they are killed, effectively enabling the female immune response to monitor its own effectiveness and to shut down when no longer needed. Elimination of this mechanism improved bacterial killing by the female immune response. This is not the first demonstration of a difference in infection outcomes between the sexes originating from differential regulation of innate immune sensing; for example, in mice, muting the inhibitory receptor CD200 resulted in greater immune activity and viral clearance, but this effect was more pronounced in female mice [40]. However, to our knowledge, this is the first case in which differential immune regulation between the sexes has been shown to result from differential degradation of microbial immune elicitors.

Stricter regulation of the Imd pathway by females suggests that immune activity may come at a greater burden to them. Uninfected wild-type females had a median survival 9.6% greater than females injected with: PBS, heat-killed *E. coli* and live *E. coli* (S1 Table). In contrast, only injection with live *E. coli* affected male survival (down 11.7% from uninfected). Because heat-killed *E. coli* are able to activate the immune response without causing mortality (shown here and [41]), these findings indicate that immune activation comes at a greater cost to females. The same trend can be seen in *PGRP-LB^Δ^* flies where male survival was only affected by injection with live *E. coli* (down 17.9% from uninfected), whereas female survival was negatively impacted equally by PBS and *E.* coli injection (down 23.9% from uninfected) (S4 Table). Together these data support the idea that the Imd response is costly (*PGRP-LB^Δ^* flies were more negatively impacted than wild-type) and that its activity poses a greater burden to females. An alternative idea is that the energy demand of *E. coli* infection in *PGRP-LB^Δ^* flies (as indicated through the decrease in both circulating and stored carbohydrate) was pathogen-derived rather than immune. Bacteria have been shown to utilize host resources during infection [15,42,43] and while this would be surprising in this infection as bacterial numbers were declining (and were also lower than that observed in wild-type flies, in which carbohydrate loss was absent – and *PGRP-LB^Δ^* flies, it remains a possibility. Indeed, the depletion of circulating sugars and glycogen in *PGRP-LB^Δ^* flies supports a model of pathogen-derived glycogenolysis [43].

Elimination of *PGRP-LB* resulted in elevated antimicrobial peptide production and thus, unsurprisingly, *PGRP-LB^Δ^* flies had roughly 1/10 the bacterial load of wild-type over the first 6h post-infection (Fig. 1E, Fig. 4D). The absence of triglyceride loss in these animals, associated with increased immune responses and reduced microbial loads, suggests that in this infection triglyceride is lost because of direct pathogen effects. We have recently shown that when flies infected with the Gram-negative pathogen *Francisella novicida* were treated with antibiotics to keep bacterial numbers low, they did not exhibit infection-driven metabolic shifts (including triglyceride loss). In contrast, when bacterial numbers increased (still in the presence of antibiotic treatment), metabolic shifts during infection were again observed, suggesting that these changes were associated with bacterial load rather than being a direct effect of the antibiotics on metabolism [33]. This idea of a ‘tipping point’ for infection pathology is likely to be a fruitful area for future inquiry.

The immune response, as we normally envision it, includes responses to infection that protect the host by killing pathogens or restricting their growth (resistance). In contrast, tolerance is defined as the ability to maintain health during infection. Experimentally, a more tolerant host is one that remains healthy longer at a given pathogen load [44,45]. Recent years have seen increasing interest in tolerance, driven in part by the idea of improving tolerance as a therapeutic approach to infection. However, despite the large body of theory surrounding tolerance, the ability to detect tolerant phenotypes [46], and the identification of tolerance-associated genes [31,44,47], we still know very little about the fundamental mechanisms of tolerance. It has previously been shown that *PGRP*-*LB* contributes to infection tolerance [38]; we show that this activity is in fact sexually dimorphic. We show that phenomenological differences in tolerance between the sexes can be used to identify fundamental mechanisms of infection tolerance and that the sex-specific regulation of inhibitors of immune signalling can underlie strong, complex differences in immune dynamics between the sexes.

## Methods

### *Drosophila* genetics and culture

*w*^1118^ flies and *w*^1118^; *imd*^10191^ were used as wild-type and Imd pathway mutants, respectively. The *imd*^10191^ line carries a 26-nucleotide deletion that frameshifts the *imd* protein at amino acid 179, which is the beginning of the death domain [48]. *PGRP-LB*^Δ^ mutant lines used were obtained from the Bloomington Stock Center and have been previously described [49]. Both mutants were placed on our *w*^1118^ genetic background using isogenic balancer chromosome lines. Flies were maintained on a sugar-yeast diet (10% w/v autolysed brewer’s yeast, 8% fructose, 2% polenta, 0.8% agar, supplemented with propionic acid and nipagin) at 25°C.

### *Drosophila* infection

For all experiments, flies were collected within 24h following eclosion and kept in same-sex vials for 5 - 7 days in groups of 20. Thus, all experiments were conducted on flies between 5 and 8 days old. Injections were carried out using a pulled-glass capillary needle and a Picospritzer injector system (Parker, New Hampshire, US). Following injection flies were kept at 29°C. Bacteria were grown from single colonies overnight at 37°C shaking. Each fly was injected with 50 nanolitres of *E. coli* suspended in PBS (OD_600_ = 1.0 ~100,000 bacteria). Following re-suspension in PBS, a subset of bacteria designated for the ‘heat-killed’ treatment was incubated for 1h at 65°C. Sterile PBS was used as a wounding control. A subset of *imd* flies were pre-injected with 0.2μm latex beads, FluoSpheres, Carboxylate-Modified Microspheres (Invitrogen) to inhibit phagocytosis as previously described [30,48]. Briefly, beads were washed 3x in sterile PBS and resuspended in PBS at one fourth of the original volume of the bead stock. Flies were injected with 50nL of bead-PBS solution or PBS alone, left for 16h, and then injected with PBS or *E. coli*.

### Survival assays

Survival experiments were performed at 29°C with 15-20 flies/vial. Survival was monitored daily and flies were tipped into fresh vials every 4 days.

### Bacterial quantification

For each sample, 1 fly was homogenised in 100μl of sterile ddH_2_O. Homogenates were serially diluted and plated onto LB agar plates where they incubated for 16-18h. Following incubation, the number of individual bacterial colonies observed on each plate was quantified and back-calculated to determine the number of CFUs present in each fly.

### Gene expression – Quantitative reverse transcription PCR

For each sample, 3 flies were homogenised in 100μl of the single-step RNA isolation reagent TRI Reagent (Sigma), followed by a chloroform extraction and precipitation in isopropanol. The resultant pellet was then washed with 70% ethanol. Pellets were resuspended and subject to DNase treatment. Revertaid M-MuLV reverse transcriptase and random hexamers (Thermo Scientific) were used to carry out cDNA synthesis. 5μl of each cDNA sample was put into a ‘neat’ standards tube; this tube was later used to generate standards which were used to generate a standard curve for each gene. Each cDNA sample was diluted and this diluted sample used for analysis.

We used Sensimix with SYBR Green no-ROX (Bioline) or qPCRBIO SyGreen Mix Separate-ROX (PCR Biosystems) for qRT-PCR. Reactions were run on a Corbett Rotor-Gene 6000 with cycling conditions as follows: Hold 95°C for 10 min, then 45 cycles of 95°C for 15s, 59°C for 30s, 72°C for 30s, followed by a melting curve. Gene expression was calculated based on the standard curve generated during each run, normalized to the value of our housekeeping gene, *Rpl1*. Samples from PBS and infected treatments were then divided by the mean value of their uninfected controls to generate expression values relative to uninfected flies.

All gene expression experiments were performed at least twice, with three or more biological replicates per experiment.

### Measurement of triglycerides

Triglycerides were measured using thin layer chromatography (TLC) assays as described elsewhere [50]. Briefly, each sample consisted of 10 flies; flies were placed in microcentrifuge tubes and stored at −80°C until the time of analysis. To perform the TLC assay, samples were removed from the −80°C freezer and spun down (3 min at 13,000 rpm at 4°C) in 100μl of a 3:1 (v/v) mix of chloroform and methanol. Flies were then homogenised and subject to a further ‘quick spin’. Standards were generated using lard dissolved in the same chloroform: methanol solution. We loaded 2μl of each standard and 20μl of each sample onto a silica gel glass plate (Millipore). Plates were then placed into a chamber pre-loaded with solvent (a 4:1 (v/v) mix of hexane and ethyl ether) and left to run until the solvent reached a point 1cm short of the edge of the plate. Plates were then removed from the chamber, allowed to dry, and stained with CAM solution [50]. Plates were baked at 80°C for 15-25min and imaged using a scanner. Triglyceride was quantified in Image J using the Gel Analysis tool.

### Measurement of carbohydrates (glucose + trehalose and glycogen)

Each sample consisted of 3 flies that were homogenised in 75μl of TE + 0.1% Triton X-100 (Sigma Aldrich). Samples were incubated for 20 min at 75°C and stored at −80°C. Prior to the assay, samples were incubated for 5 min at 65°C. Following incubation, 10μl from each sample was loaded into 4 wells of a 96-well plate. Each well was designated to serve as a measurement for either: control (10μl sample + 190μl H_2_0), glucose (10μl sample + 190μl glucose reagent (Sentinel Diagnostics)), trehalose (10μl sample + 190μl glucose reagent + trehalase (Sigma Aldrich)), or glycogen (10μl sample + 190μl glucose reagent + amyloglucosidase (Sigma Aldrich)). A standard curve was generated by serially diluting a glucose sample of known concentration and adding 190μl of glucose reagent to 10μl of each standard. Standards were always run at the same time and in the same plate as samples. Plates were incubated for 1.5h −3h at 37°C following which the absorbance for each well at 492 nm was determined using a plate reader.

### Respirometry

Respiration in flies was measured using a stop-flow gas-exchange system (Q-Box RP1LP Low Range Respirometer, Qubit Systems, Ontario, Canada, K7M 3L5). Eight flies from each treatment were put into an airtight glass tube and supplied with our standard fly food via a modified pipette tip. Each tube was provided with CO_2_-free air while the ‘spent’ air was concurrently flushed through the system and analysed for its CO_2_ and O_2_ content. In this way, evolved CO_2_ and consumed O_2_ were measured for each tube every ~ 44 min (the time required to go through each of the 7 vials in sequence). For most replicates of the respirometry assay, there were 2 uninfected, 2 PBS and 3 infected vials.

### Reproductive assay

Flies were collected within 7 hours of eclosion to ensure virginity. To assess fitness, immediately following injection with either PBS or *E. coli*, flies were placed into vials with uninfected competitors of the same sex and potential mates of the opposite sex. Competitor flies expressed DsRed marker eyes, this marker allowed for easy identification of offspring resulting from focal flies - any DsRed eyed offspring were the progeny of competitor flies. Flies were allowed to mate for 12h as this interval exceeds the time required for flies to significantly reduce the number of – and by some reports, clear - *E*. *coli*, thus, allowing us to observe fitness throughout the infection. In one block, *E. coli* reproductive assays were left for 24h, we have included these data as number of offspring produced did not differ from the shorter assay, possibly because females do not lay many eggs overnight. After the mating period, flies were discarded and vials were left for 14 days to allow resultant offspring time to develop and eclose.

### Statistical analysis

Data were analysed in R Studio with R version 3.5.1 [51]. Survival data were initially analysed using Cox proportional hazards models; we then used Log-Rank tests for pairwise comparisons. We ran a GLM of reproductive success by sex and infection treatment; then, using only those matings resulting in offspring, we performed a GLM on number of offspring produced by sex and infection treatment. For all other assays, we first tested for normality of data which dictated whether an ANOVA, t-test, Kruskal – Wallis analysis of variance, or Mann-Whitney U test was used to calculate differences between treatments with sex and infection status as factors. All initial models included experimental replicate as a factor and failed to observe an effect. When appropriate, we performed post-hoc Tukey or Dunn analyses to identify specific differences between treatments. All assays consist of 2-4 replicates.

## Acknowledgements

Members of the Dionne lab, the Imperial SK fly lab and D. Duneau provided useful feedback on the manuscript. This work was supported by MRC Research Grant (MR/R00997X/1) and Wellcome Trust Investigator Award 207467/Z/17/Z.

## Competing interests

No competing interests.

## Supporting data

**Figure S1.**
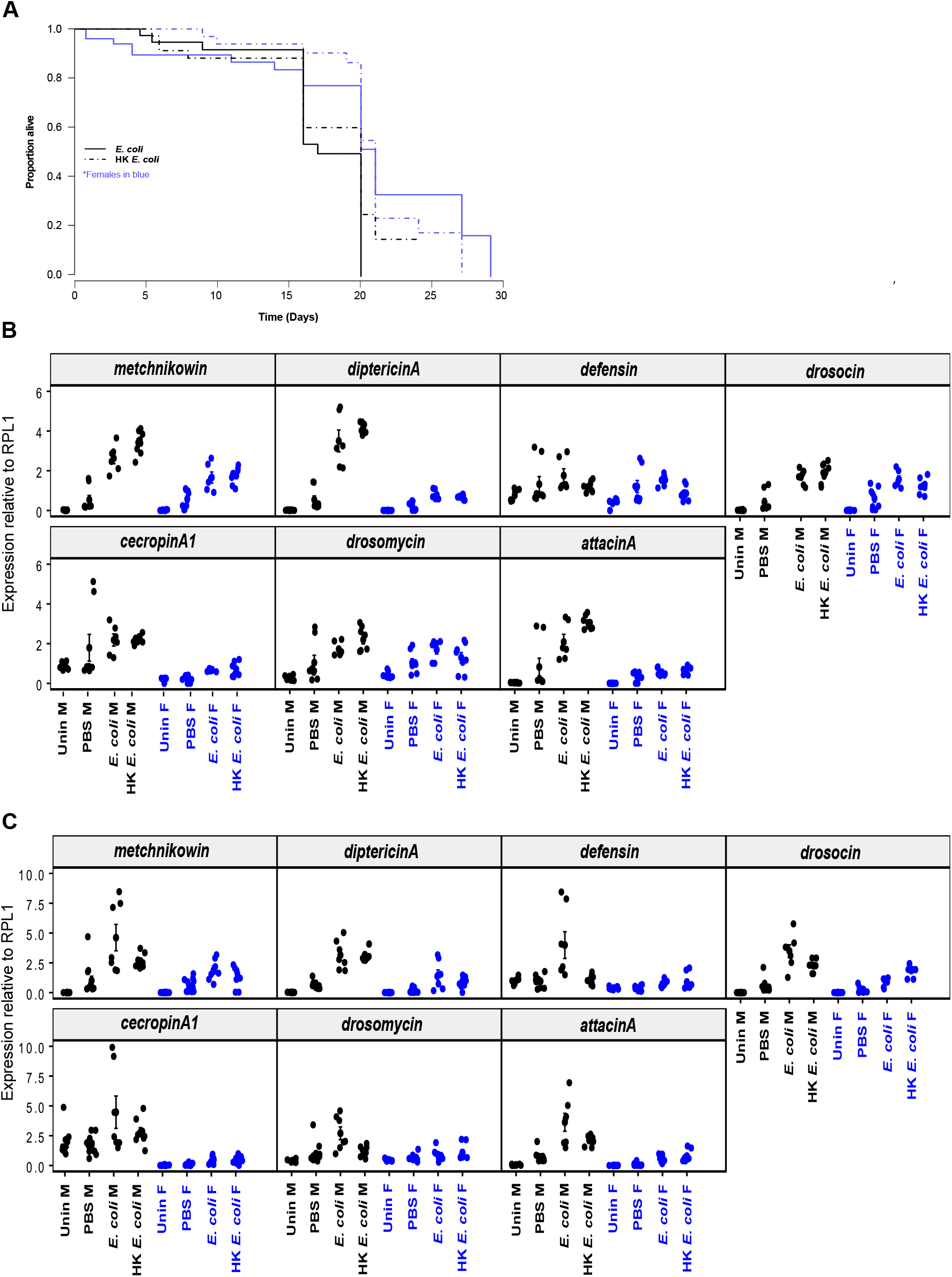
**(A)** Survival of wild-type flies infected with live and heat-killed (HK) *E. coli*. Males and females are represented by black and blue tracings, respectively. HK bacteria were incubated for 1h at 65 C. Live *E. coli* data are replotted from figure 1a. Antimicrobial peptide expression **(B)** 3h and **(C)** 6h following *E. coli* injection. All genes were standardized to the housekeeping gene ribosomal protein 1. Data are presented in arbitrary units. Markers represent individual data points. Bars indicate SE. All assays were performed twice, each repeat included 3-4 biological replicates/treatment consisting of 3 flies each.

**Figure S2.**
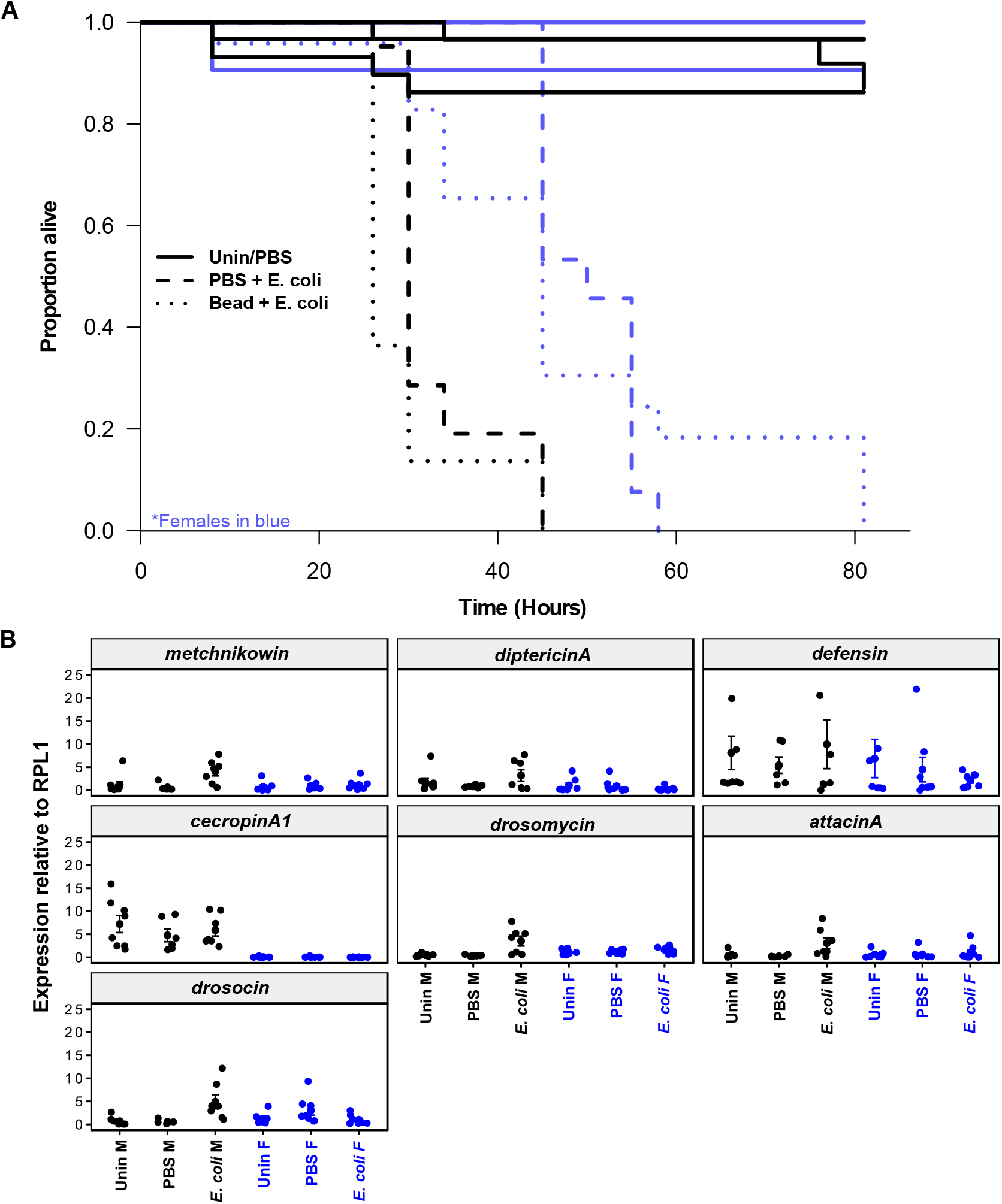
**(A)** Survival of imd mutants pre-injected with either beads or PBS prior to *E. coli*. In boxplot, median value is indicated by horizontal bars, top and bottom of boxes represent upper and lower quartiles (respectively). Whiskers indicate maximum and minimum values. **(B)** Antimicrobial peptide expression 6h following *E. coli* injection in imd mutant flies. All genes were standardized to the housekeeping gene ribosomal protein 1. Data are presented in arbitrary units. Markers represent individual data points. Bars indicate SE. All assays were performed twice, each repeat included 3-4 biological replicates/treatment consisting of 3 flies each.

**Figure S3.**
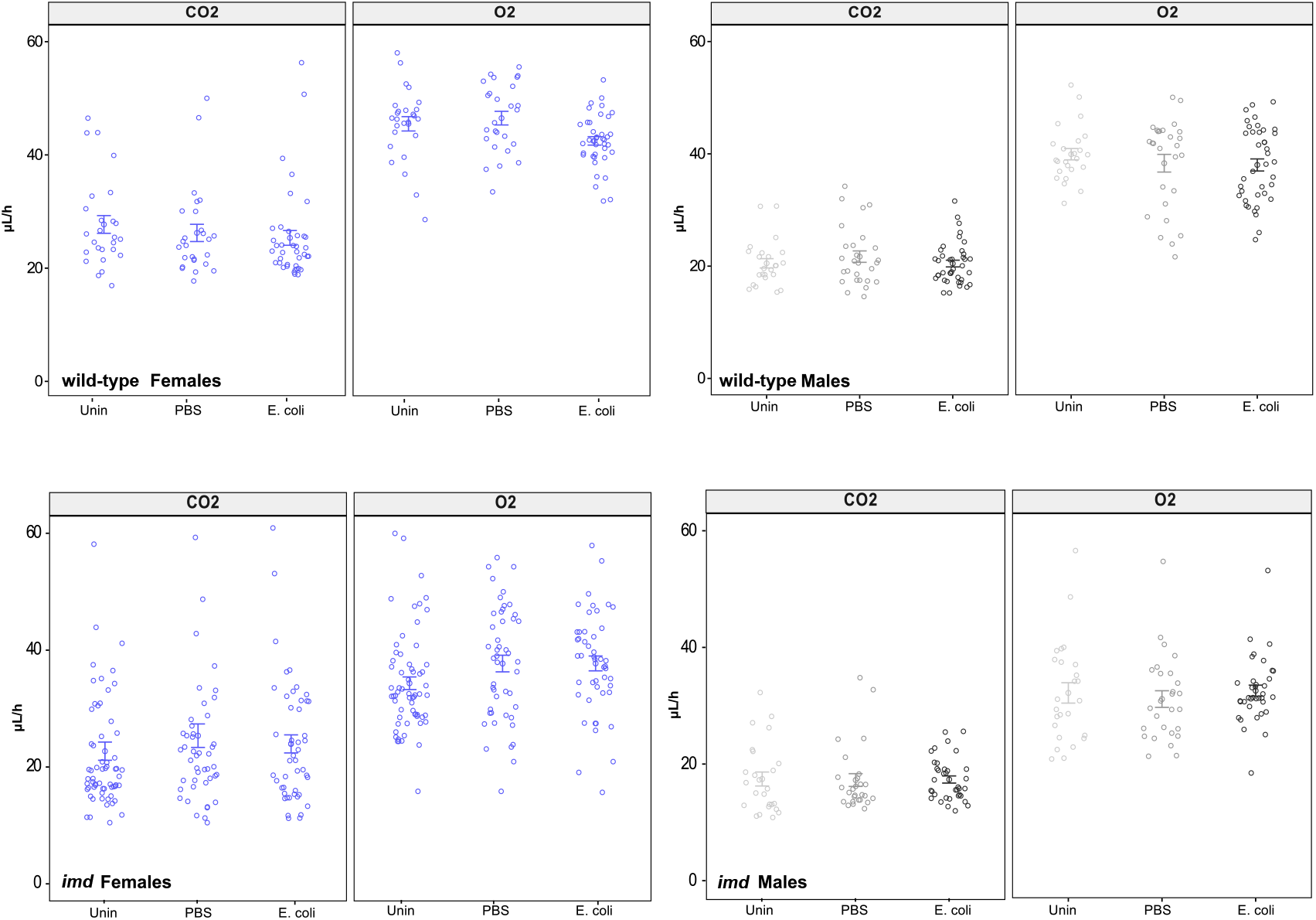
Respiration of infected males and females. Respiration was measured for six hours following infection. Markers represent one vial consisting of 8 flies. Bars indicate SE. All assays were repeated 3 or 4 times with 2 or 3 samples/treatment.

**Figure S4.**
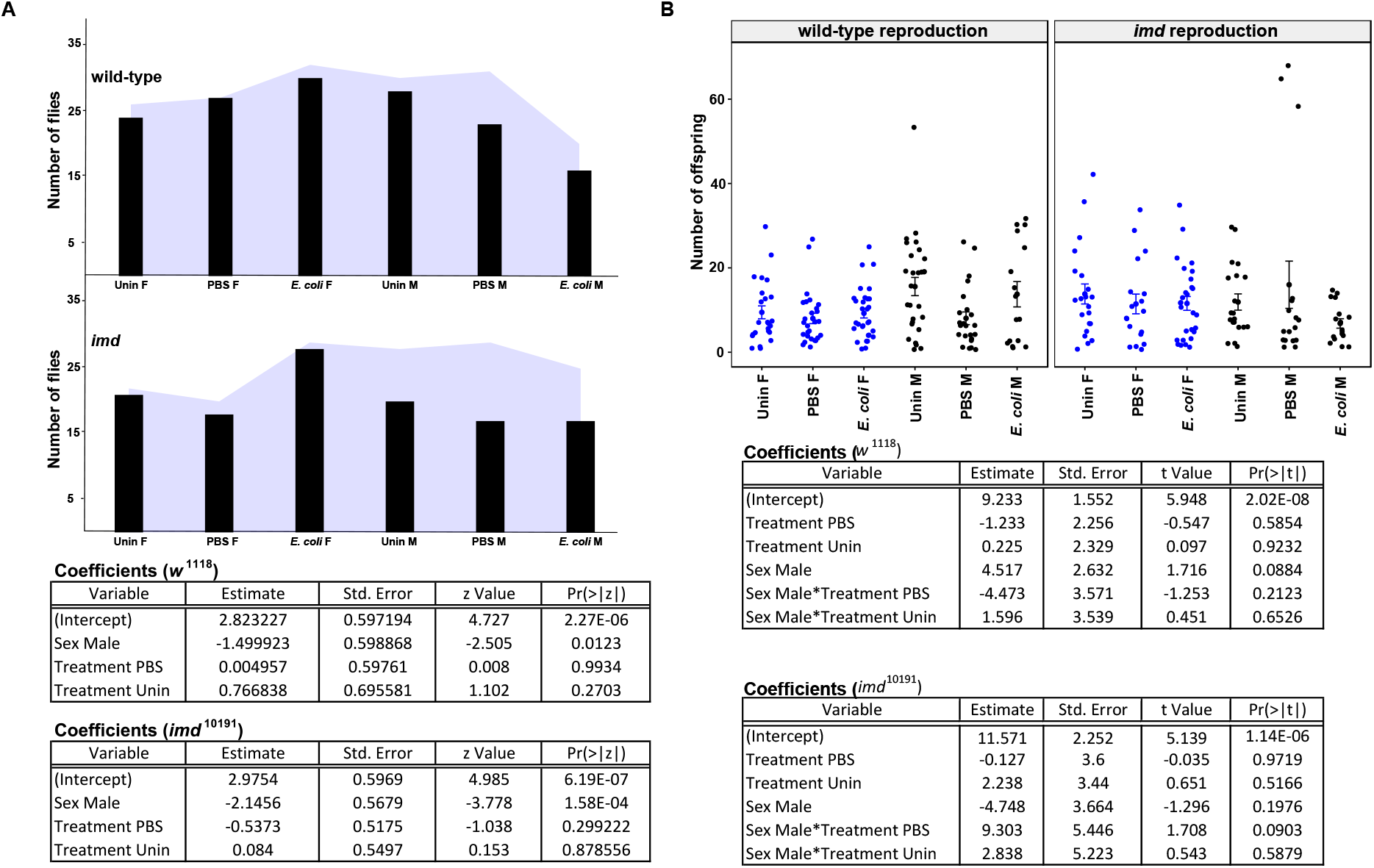
Reproductive success during *E. coli* infection. **(A)** proportion of flies from each treatment that successfully mated. Blue shaded area represents the total number of flies put into mating assay;black bars show the number of flies that produced at least one (1) adult offspring. **(B)** Number of adult offspring resulting from 10/12h mating assays. Large markers indicate means while smaller circles represent individual assays. Bars indicate SE. Experiments were performed at least twice, n= 8-15 biological replicates each. Output from GLM models are shown.

**Figure S5.**
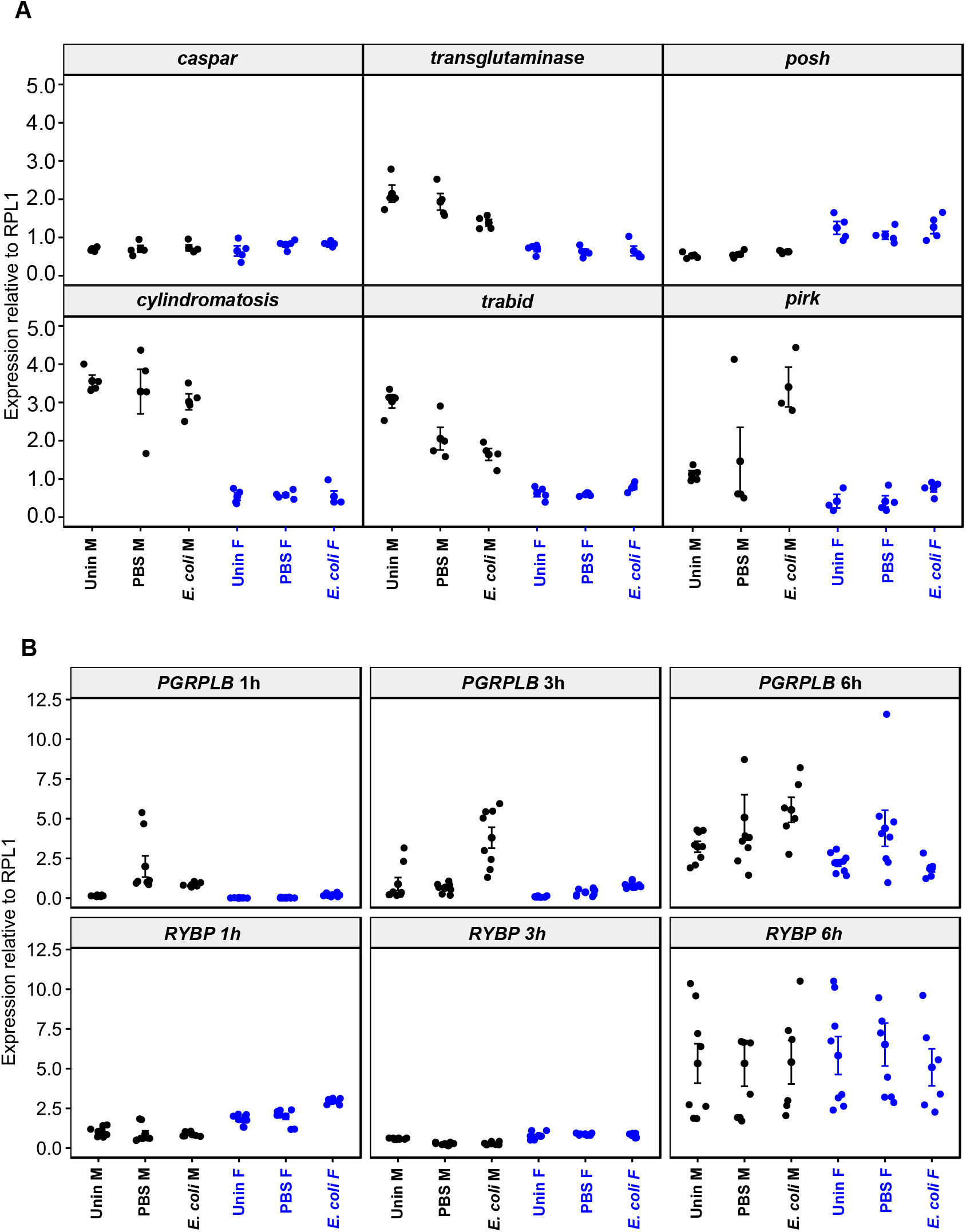
Regulators of the Imd pathway. **(A)** Expression of known Imd pathway regulators 6h following *E. coli* injection and **(B)** a time course of regulators *PGRP-LB* and *RYBP*. Genes were standardized to the housekeeping gene ribosomal protein 1. Data are presented in arbitrary units. Markers represent individual data points. Bars indicate SE. We used 3 or 4 biological replicates/gene, consisting of 3 flies. Time course performed twice.

**Figure S6.**
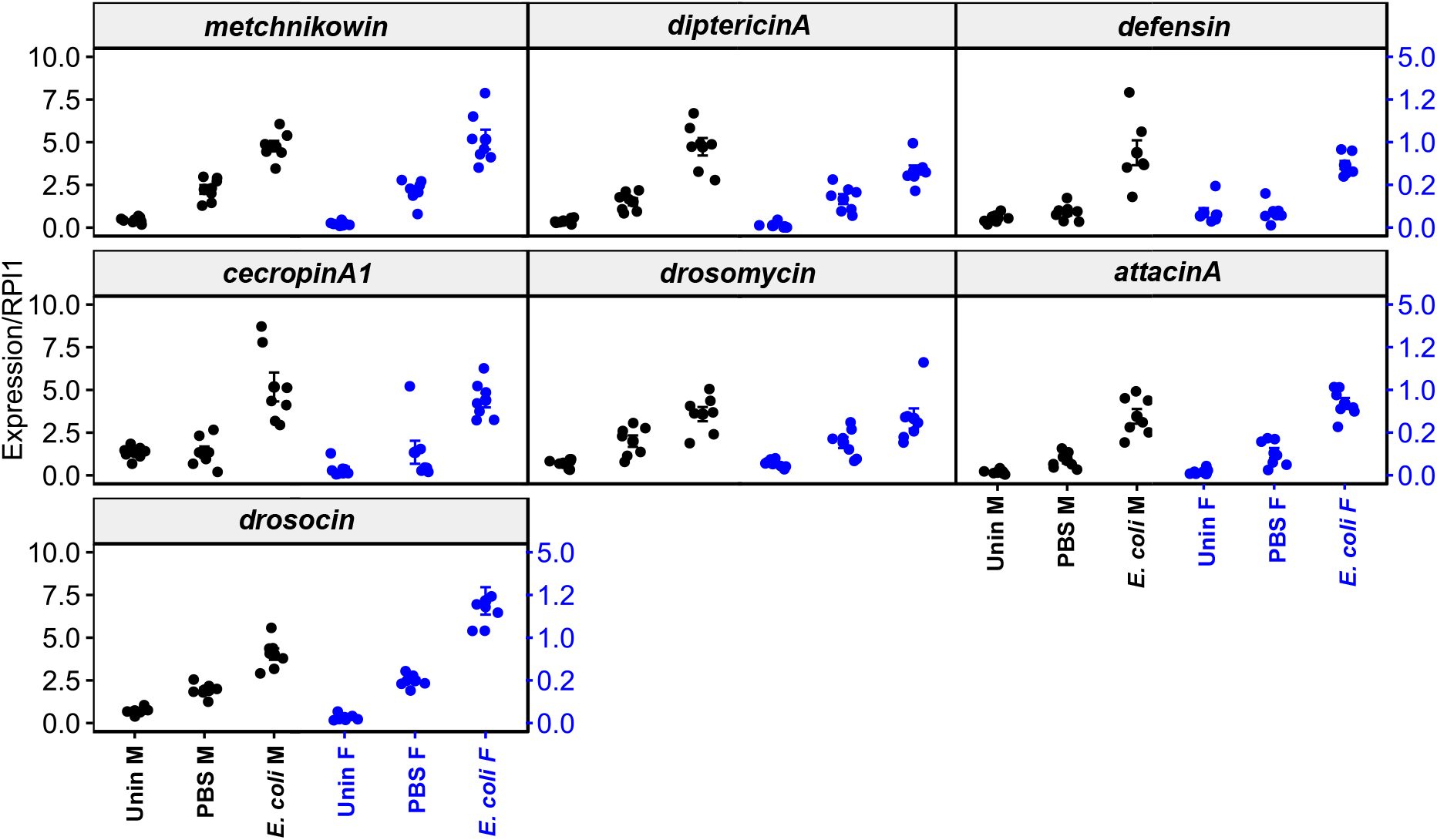
Antimicrobial peptide expression 6h following *E. coli* injection in *PGRP-LB*Δ flies. All genes were standardized to the housekeeping gene ribosomal protein 1. Data are presented in arbitrary units. Markers represent individual data points. Bars indicate SE. All assays were performed twice, each repeat included 3-4 biological replicates/treatment consisting of 3 flies each.

**Supplementary table 1:** wild-type median survival (days)

| Treatment | Median survival | upper CI | lower CI | % change |
| --- | --- | --- | --- | --- |
| Uninfected M | 18.8 | 19.8 | 15.8 | - |
| PBS M | 19.8 | 20.8 | 18.8 | +5.3 |
| HK *E. coli* M | 19.8 | 20.8 | 15.8 | +5.3 |
| *E. coli* M | 16.8 | NA | 15.8 | -11.7 |
| Uninfected F | 22.8 | 26.8 | 19.8 | - |
| PBS F | 20.8 | 25.8 | 20.8 | -9.6 |
| HK *E. coli* F | 20.8 | 20.8 | 19.8 | -9.6 |
| *E. coli* F | 20.8 | NA | 19.8 | -9.6 |

\*HK - heat-killed; M - males; F - females;
% change from uninfected control

**Supplementary table 2:**
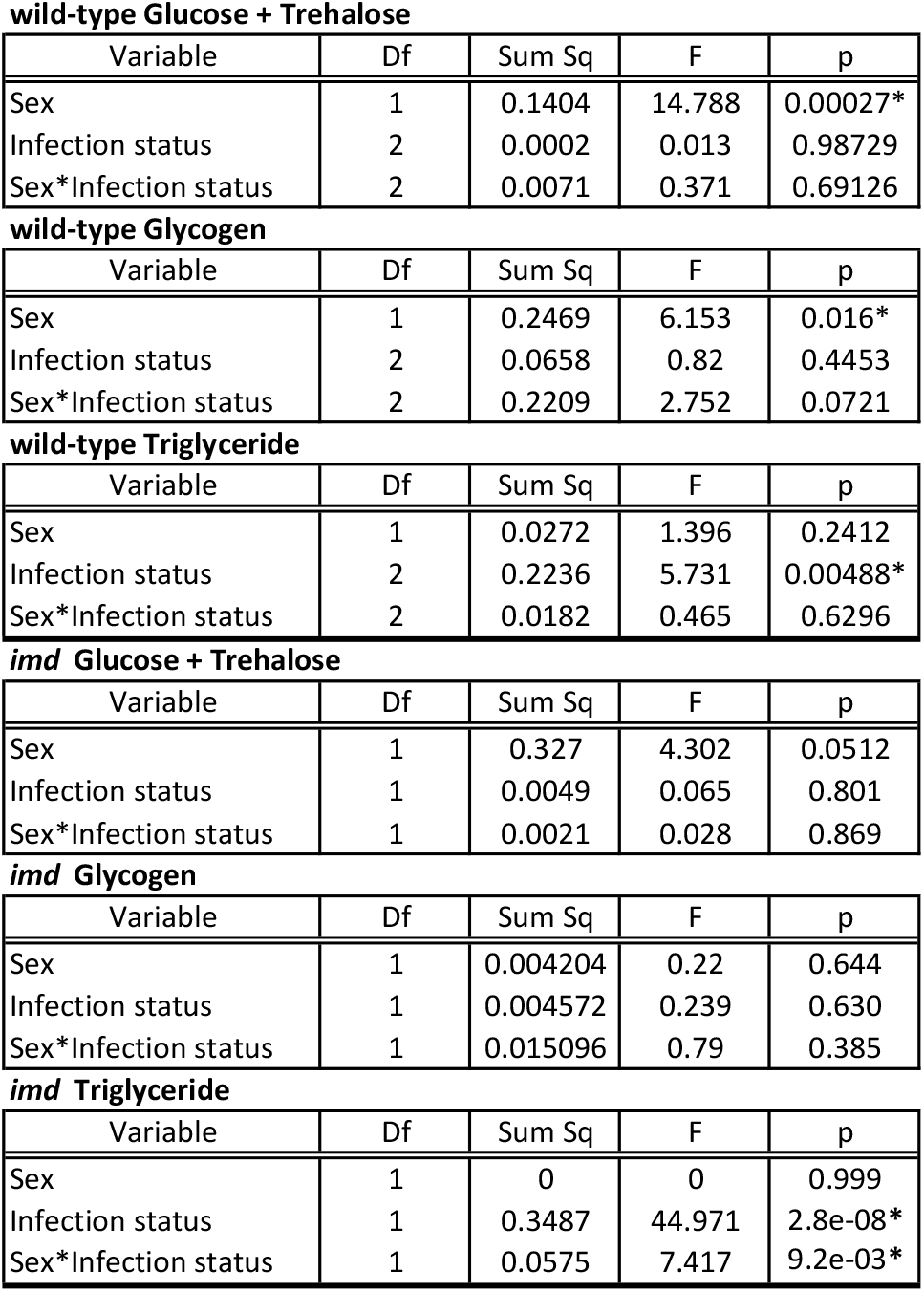
Metabolic statistics.

**wild-type Glucose + Trehalose**
| Variable | Df | Sum Sq | F | p |
| --- | --- | --- | --- | --- |
| Sex | 1 | 0.1404 | 14.788 | 0.00027* |
| Infection status | 2 | 0.0002 | 0.013 | 0.98729 |
| Sex*Infection status | 2 | 0.0071 | 0.371 | 0.69126 |

**wild-type Glycogen**
| Variable | Df | Sum Sq | F | p |
| --- | --- | --- | --- | --- |
| Sex | 1 | 0.2469 | 6.153 | 0.016* |
| Infection status | 2 | 0.0658 | 0.82 | 0.4453 |
| Sex*Infection status | 2 | 0.2209 | 2.752 | 0.0721 |

**wild-type Triglyceride**
| Variable | Df | Sum Sq | F | p |
| --- | --- | --- | --- | --- |
| Sex | 1 | 0.0272 | 1.396 | 0.2412 |
| Infection status | 2 | 0.2236 | 5.731 | 0.00488* |
| Sex*Infection status | 2 | 0.0182 | 0.465 | 0.6296 |

**imd Glucose + Trehalose**
| Variable | Df | Sum Sq | F | p |
| --- | --- | --- | --- | --- |
| Sex | 1 | 0.327 | 4.302 | 0.0512 |
| Infection status | 1 | 0.0049 | 0.065 | 0.801 |
| Sex*Infection status | 1 | 0.0021 | 0.028 | 0.869 |

**imd Glycogen**
| Variable | Df | Sum Sq | F | p |
| --- | --- | --- | --- | --- |
| Sex | 1 | 0.004204 | 0.22 | 0.644 |
| Infection status | 1 | 0.004572 | 0.239 | 0.630 |
| Sex*Infection status | 1 | 0.015096 | 0.79 | 0.385 |

**imd Triglyceride**
| Variable | Df | Sum Sq | F | p |
| --- | --- | --- | --- | --- |
| Sex | 1 | 0 | 0 | 0.999 |
| Infection status | 1 | 0.3487 | 44.971 | 2.8e-08* |
| Sex*Infection status | 1 | 0.0575 | 7.417 | 9.2e-03* |

**Supplementary table 3:**
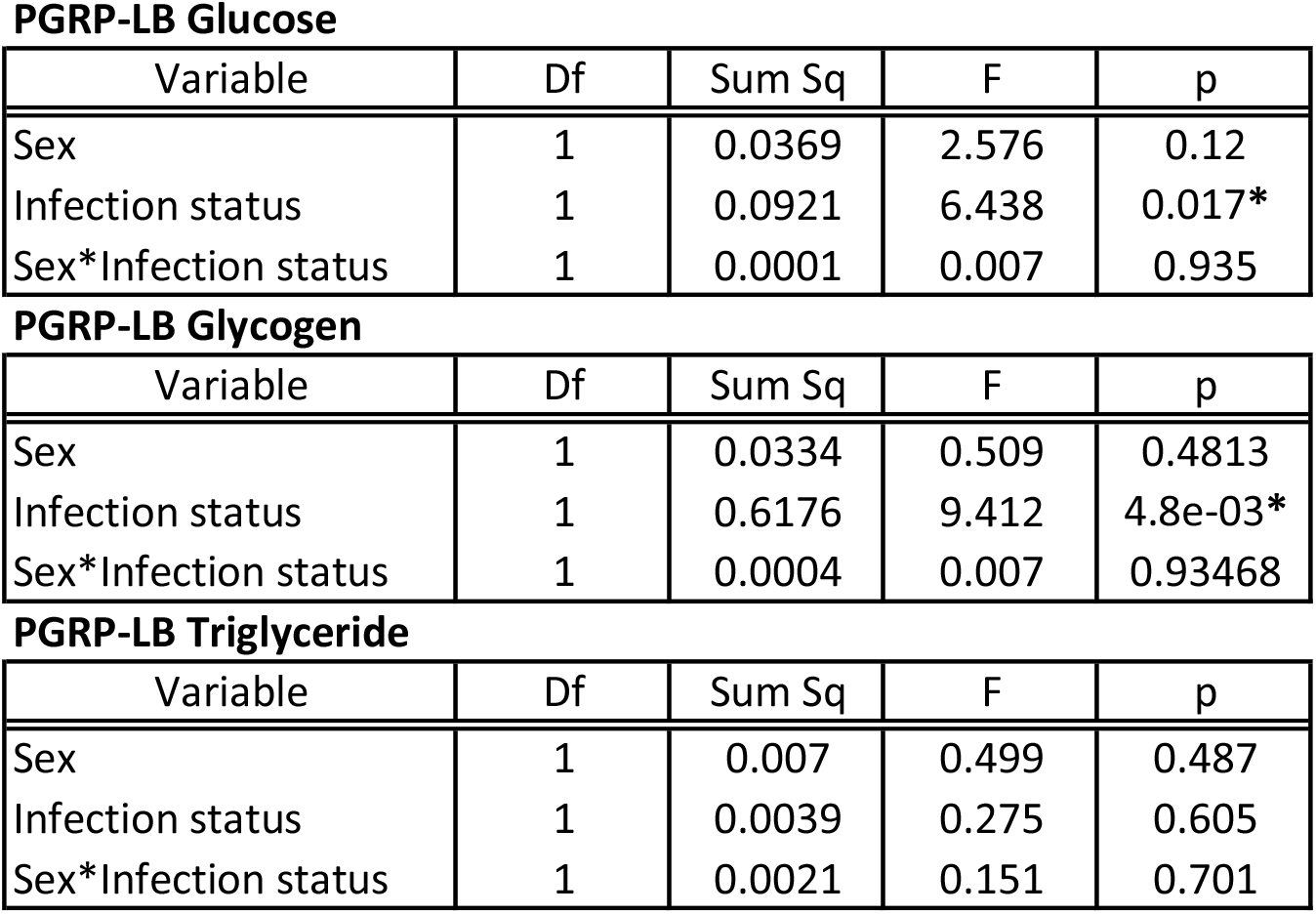
Metabolic statistics.

**Supplementary table 4:** *PGRP-LB* median survival (days)

| Treatment | Median survival | upper CI | lower CI | % change |
| --- | --- | --- | --- | --- |
| Uninfected M | 13.2 | 13.2 | 13.2 | - |
| PBS M | 13.2 | 13.2 | 11.2 | 0 |
| *E. coli* M | 11.2 | 13.2 | 10.2 | -17.9 |
| Uninfected F | 25.9 | 25.9 | 20.9 | - |
| PBS F | 20.9 | 20.9 | 17.3 | -23.9 |
| *E. coli* F | 20.9 | 20.9 | 20.9 | -23.9 |

\*M - males; F - females; % change from uninfected control

## Notes

### Competing Interest Statement

The authors have declared no competing interest.

